# Massive gene presence/absence variation in the mussel genome as an adaptive strategy: first evidence of a pan-genome in Metazoa

**DOI:** 10.1101/781377

**Authors:** Marco Gerdol, Rebeca Moreira, Fernando Cruz, Jessica Gómez-Garrido, Anna Vlasova, Umberto Rosani, Paola Venier, Miguel A. Naranjo-Ortiz, Maria Murgarella, Pablo Balseiro, André Corvelo, Leonor Frias, Marta Gut, Toni Gabaldón, Alberto Pallavicini, Carlos Canchaya, Beatriz Novoa, Tyler S. Alioto, David Posada, Antonio Figueras

## Abstract

Mussels are ecologically and economically relevant edible marine bivalves, highly invasive and resilient to biotic and abiotic stressors causing recurrent massive mortalities in other species. Here we show that the Mediterranean mussel *Mytilus galloprovincialis* has a complex pan-genomic architecture, which includes a *core* set of 45,000 genes shared by all individuals plus a surprisingly high number of *dispensable* genes (∼15,000). The latter are subject to presence/absence variation (PAV), i.e., they may be entirely missing in a given individual and, when present, they are frequently found as a single copy. The enrichment of *dispensable* genes in survival functions suggests an adaptive value for PAV, which might be the key to explain the extraordinary capabilities of adaptation and invasiveness of this species. Our study underpins a unique metazoan pan-genome architecture only previously described in prokaryotes and in a few non-metazoan eukaryotes, but that might also characterize other marine invertebrates.

**Significance statement:** In animals, intraspecific genomic diversity is generally thought to derive from relatively small-scale variants, such as single nucleotide polymorphisms, small indels, duplications, inversions and translocations. On the other hand, large-scale structural variations which involve the loss of genomic regions encoding protein-coding genes in some individuals (i.e. presence/absence variation, PAV) have been so far only described in bacteria and, occasionally, in plants and fungi. Here we report the first evidence of a pan-genome in the animal kingdom, revealing that 25% of the genes of the Mediterranean mussel are subject to PAV. We show that this unique feature might have an adaptive value, due to the involvement of dispensable genes in functions related with defense and survival.

## Background

The Mediterranean mussel *Mytilus galloprovincialis* Lamarck, 1819 (Bivalvia, Mytilida), a member of the *M. edulis* species complex, is an edible cosmopolitan bivalve mollusk with a high socio-economic impact. This shellfish has been consumed by humans since 6000 BC, and its global production currently exceeds 400 thousand tons per year (1). Due to its invasive nature, this species has spread far beyond its native range, and it is considered a worldwide threat for autochthonous bivalve populations (2). Upon the release of gametes into the open water, and after fertilization, larvae can travel long distances carried by the oceanic currents (3). Metamorphosis takes place during planktonic life, which ends after one to two months – depending on water temperature and food availability– with the settlement of juveniles and the start of the sessile adult life. Mussel beds are therefore usually composed by genetically heterogeneous individuals derived from large, randomly-mating populations of different geographical origin. As filter-feeders, mussels are constantly exposed to a wide range of potentially pathogenic microorganisms, biotoxins and anthropogenic pollutants. However, they display a remarkable resilience to stress and infection, can evolve novel traits in response to predation within a few generations (4) and are capable of significant bioaccumulation (5), without experimenting the massive mortalities seen in other farmed bivalves (6, 7). The molecular mechanisms underpinning this invasiveness and resilience are poorly understood, and it is unclear whether the genomic complexity of this species (8) represents a key factor in sustaining its extraordinary capability of adaptation.

As a matter of fact, bivalve genomes are generally complex, relatively large and highly heterozygous, harboring numerous mobile elements (9–13). These factors have proved to be particularly challenging in previous assembly efforts in *M. galloprovincialis*, resulting in extremely fragmented and poorly contiguous genome sequences (8, 14). In order to carry out an in-depth investigation of intraspecific genomic divergence in this mussel, we built and improved genome assembly from a single mussel (*Lola*) before resequencing the genome of 14 individuals from separate populations in Spain and Italy. Surprisingly, we found that more than one third of the reference assembly comprises regions that are just found in one of the two homologous chromosomes of *Lola.* Moreover, these same regions were often entirely missing in the resequenced individuals. Our analyses revealed that these regions contain approximately 15,000 *dispensable* protein-coding genes, subject to presence/absence variation (PAV), in contrast with *core* genes, which are invariably present in all individuals. Remarkably, most *dispensable* genes belonged to *Mytilus*-specific expanded families, and were enriched in functions related with protein/carbohydrate recognition and survival, suggesting a potential role of PAV in the evolutionary success of this cosmopolitan marine invertebrate. This unexpected discovery represents the first evidence of widespread PAV in a metazoan, extending the concept of pan-genome, so far only considered as a relevant phenomenon in viruses, prokaryotes and a few fungi and plants (15–22), to the animal kingdom.

### An overview of the mussel reference genome

We assembled the reference genome using a combination of Illumina short-reads from paired end, mate-pair and fosmid libraries, and long reads generated with the single-molecule real-time (SMRT) Pacific Biosciences technology. We followed a multi-step hierarchical assembly strategy (**Supplementary Data Note 1**), specifically designed to accurately remove uncollapsed alleles and homologous haplotype stretches, that resulted in a 1.28 Gb genome, of slightly smaller size compared to cytogenetic estimates (23), but of higher quality and contiguity compared to previous attempts (8, 14) (10,577 scaffolds; contig N50 = 71.42 Kb; scaffold N50 = 207.64 Kb). Although this genome shares some typical features of other bivalves, such as a low GC content (∼32%) and a widespread presence of repeats (∼43% of the assembly), it has a significantly higher number of protein-coding genes than oysters and scallops (60,338) (9, 11) (**Supplementary Data Note 2-4**). The reconstruction of the evolutionary relationships among *M. galloprovincialis* and 15 selected lophotrochozoan species (24–27), followed by a phylostratigraphic analyses (28), revealed that this large gene repertoire is the result of multiple lineage-specific duplication events that took place after the split between *Mytilus* and the rest of Mytilida (**Figure 1, Supplementary Data Note 5**). Although most mussel genes were functionally annotated based on sequence similarity (56.17%), and were supported by transcriptomic evidence (78.70%), a non-negligible number of genes (more than 5,000) have been recently acquired, i.e., they are taxonomically restricted to the *Mytilus* lineage (**Supplementary Data Note 17**) (29).

**Figure 1:**
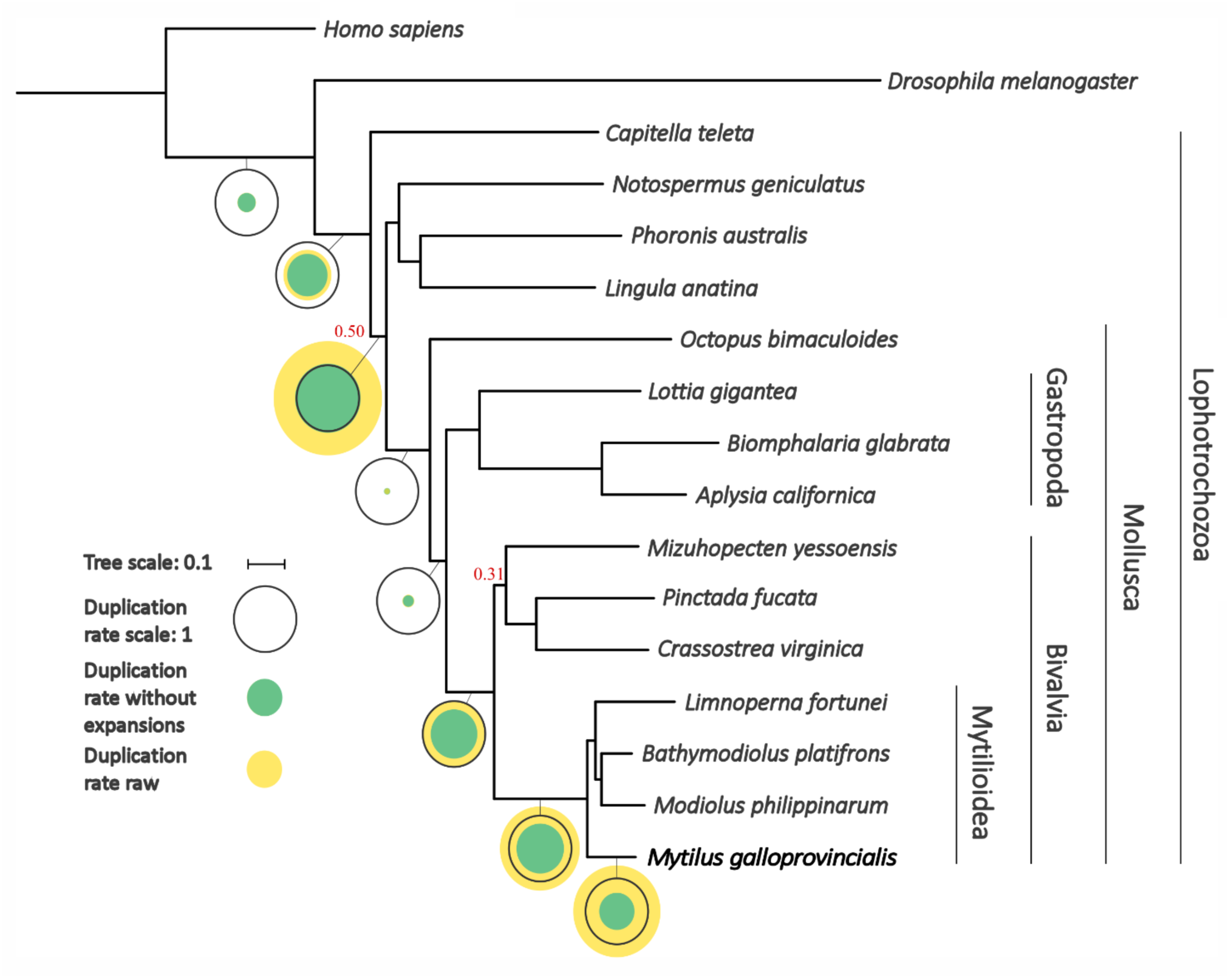
Species tree obtained from the concatenation of 177 widespread single-gene families. In bold we indicate the genome sequenced in this study. All branches were very well supported (aLRT > 0.99) except those with a number in red. The tree topology identifies Bivalvia as sister branch to Gastropoda, with both molluscan classes forming a clade sister to Cephalopoda. Mollusca appears as sister branch of a clade containing Phoronida (*Phoronis*), Nemertea (*Notospermus*) and Brachiopoda (*Lingula*) with low support (0.50). Circles represent duplication rates (genome-wide average number of duplications per gene). Yellow circles represent the estimated duplication rates before removing large expansions consisting of more than 20 paralogs appearing in a single node. Green circles represent the duplication rates obtained after the removal of such events. Black circles serve as a scale and correspond to a duplication rate of 1.

### A genome with widespread hemizygosity

Consistently with previous reports from other mytilid species (12, 13), the mussel genome is highly heterozygous. The contribution of heterozygosity to the overall intraspecific genomic variation was estimated in *Lola* and in the resequenced genomes by analyzing common regions in all individuals, in order to overcome the limitations of *k-mer*-based methods. The average heterozygosity rate observed across individuals was 1.73±0.24%, indicating that the mussel genome harbors a very high density of single nucleotide polymorphisms (SNPs), 12-22 fold higher than the human genome (30–32) (**Supplementary Data Note 6**).

However, while this high heterozygosity rate may seem rather large when compared to other animal species, it does not appear to be the main source of intraspecific genomic diversity in *M. galloprovincialis*. Indeed, in spite of the assembly strategy we adopted, the reference genome assembly still contained a high fraction (36.78%) of sequence with very low coverage. The bimodal distribution of the read coverage in *Lola* (**Figure 2B**) clearly shows that such regions are found in a *hemizygous* state, i.e., they are present in only one of the two homologous chromosomes. This mirrors the situation previously described in other metazoans with high intraspecific genome diversity, such as the roundworm *Caenorhabditis brenneri* and the ascidian *Ciona savignyi*, which have genomes characterized by significant structural variations (SV) and frequent polymorphic indels (33, 34).

**Figure 2:**
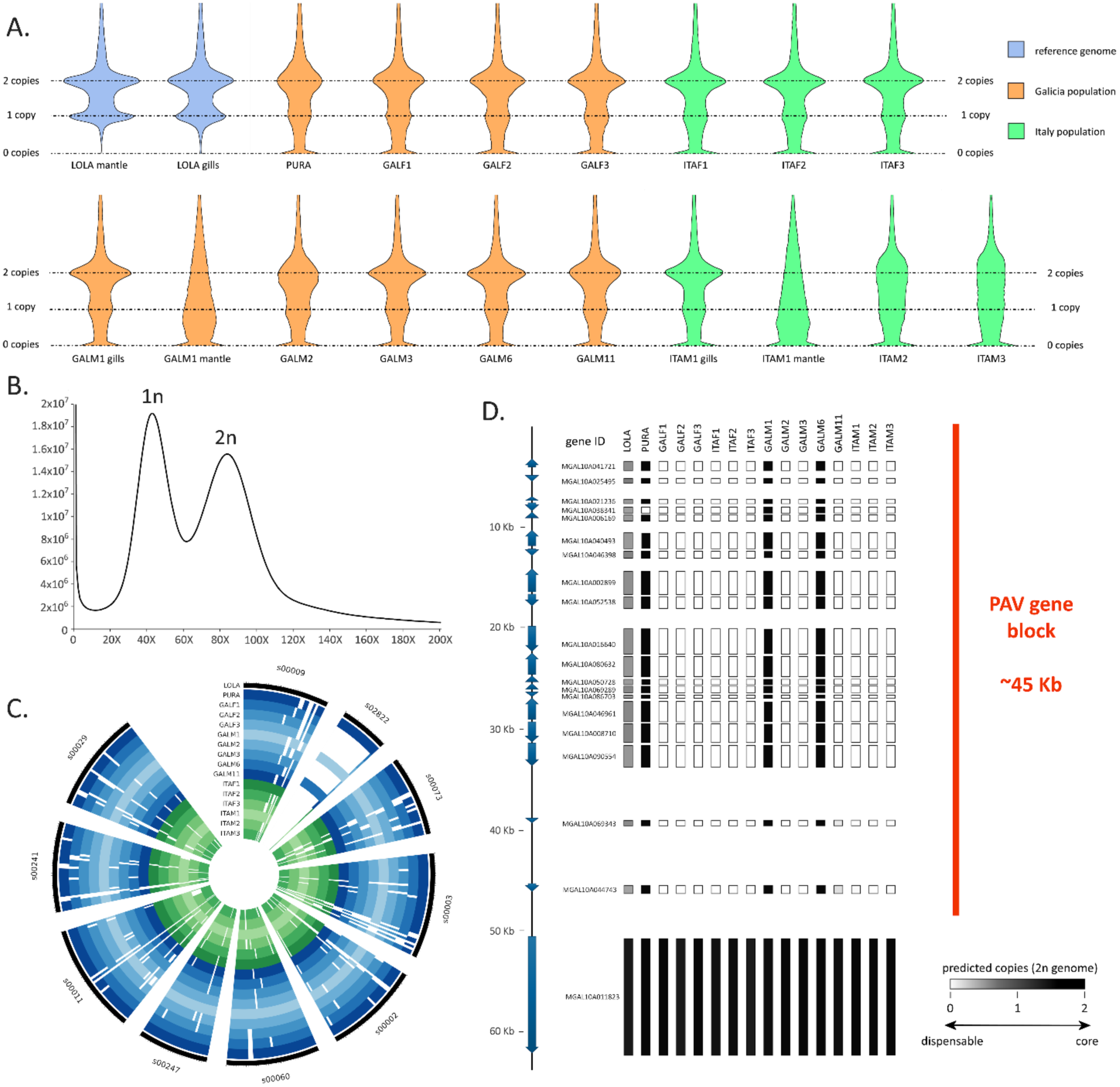
Panel A: violin plots displaying the “per gene” coverage of the protein-coding genes annotated in the mussel genomes, normalized on the expected haploid mussel genome size, calculated based on the mapping of Illumina PE libraries, obtained either from the mantle or gills tissue. Panel B: summary of the “per base” coverage of the *Lola* assembly (evaluated by the mapping of Illumina PE reads obtained from the mantle tissue). Two main peaks of coverage are clearly visible, corresponding to the haploid (1n, ∼41.5X) and diploid (2n, ∼83X) peaks of coverage based on genome size estimates. The peak located at 0 indicates approximately 70 Mb of genome assembly which did not achieve any mapping based on Q >= 60. Panel C: circus plot summarizing the presence/absence variability phenomenon on the nine longest genome scaffolds, plus the scaffold 02822, which contains a large fraction of *dispensable* genes. Italian and Galician mussel genomes are indicated with green and blue shades of color, respectively. Full and empty circles indicate present and absent genes, respectively. Panel D: Detail of presence/absence variation in the genomic scaffold 02822; note that this scaffold contains a single *core* gene and a large (about 45 Kb) block of *dispensable* genes, which are only found in four out of the analyzed genomes (including *Lola*, where the gene block is present in a single copy only).

### Massive gene presence/absence variation

Remarkably, the hemizygous fraction of the mussel genome does not only include non-coding elements and intergenic regions, but also contains a large number of protein-coding genes, which account for nearly one third of the 60,338 genes annotated in the reference genome (**Supplementary Data Note 8** and **10**). Even more surprisingly, our analyses revealed that 14,820 protein-coding genes (24.25% of the total) were entirely missing in at least one of the resequenced genomes (**Figure 2A-C**). Gene presence/absence variation (PAV) is a well-known phenomenon in prokaryotes and in a small number of eukaryotes (i.e., cultivated crops, fungi and unicellular marine algae) (15–18, 21, 22, 35), but has never been reported before on such a large scale in metazoans. Unlike the 45,518 *core* genes that are found in two copies in in *Lola* and in all the resequenced genomes, the 14,820 genes subject to PAV are *dispensable*, i.e., they can be present in either one, two or in neither of the two homologous chromosomes of the different mussels analyzed. Several additional observations are consistent with the tight association between *dispensable* genes and hemizygosity. Indeed, most of the genes present in a single copy in *Lola* were either present in a single copy or absent in the resequenced genomes (58.50% and 23.23% on average, respectively). Moreover, the overwhelming majority (98.05%) of the genes present in two copies in *Lola* were present in all the resequenced genomes, in 85.46% of cases with two copies. Finally, the significant contiguity observed for some *dispensable* genes indicates that they are contained in large (up to 30 kb) hemizygous genomic regions (**Figure 2D, Supplementary Data Note 14**). We ruled out the possibility that our observations resulted from biases related to library preparation, sequencing or bioinformatics analysis. This was achieved by: (i) resequencing *Lola*’s genome from a different tissue (gills), which fully confirmed the results obtained with the first round of sequencing; (ii) confirming the observed PAV patterns with PCRs carried out on all resequenced mussels, targeting twelve *dispensable* genes plus five *core* genes (**Figure 3A, Supplementary Data Note 11**).

**Figure 3:**
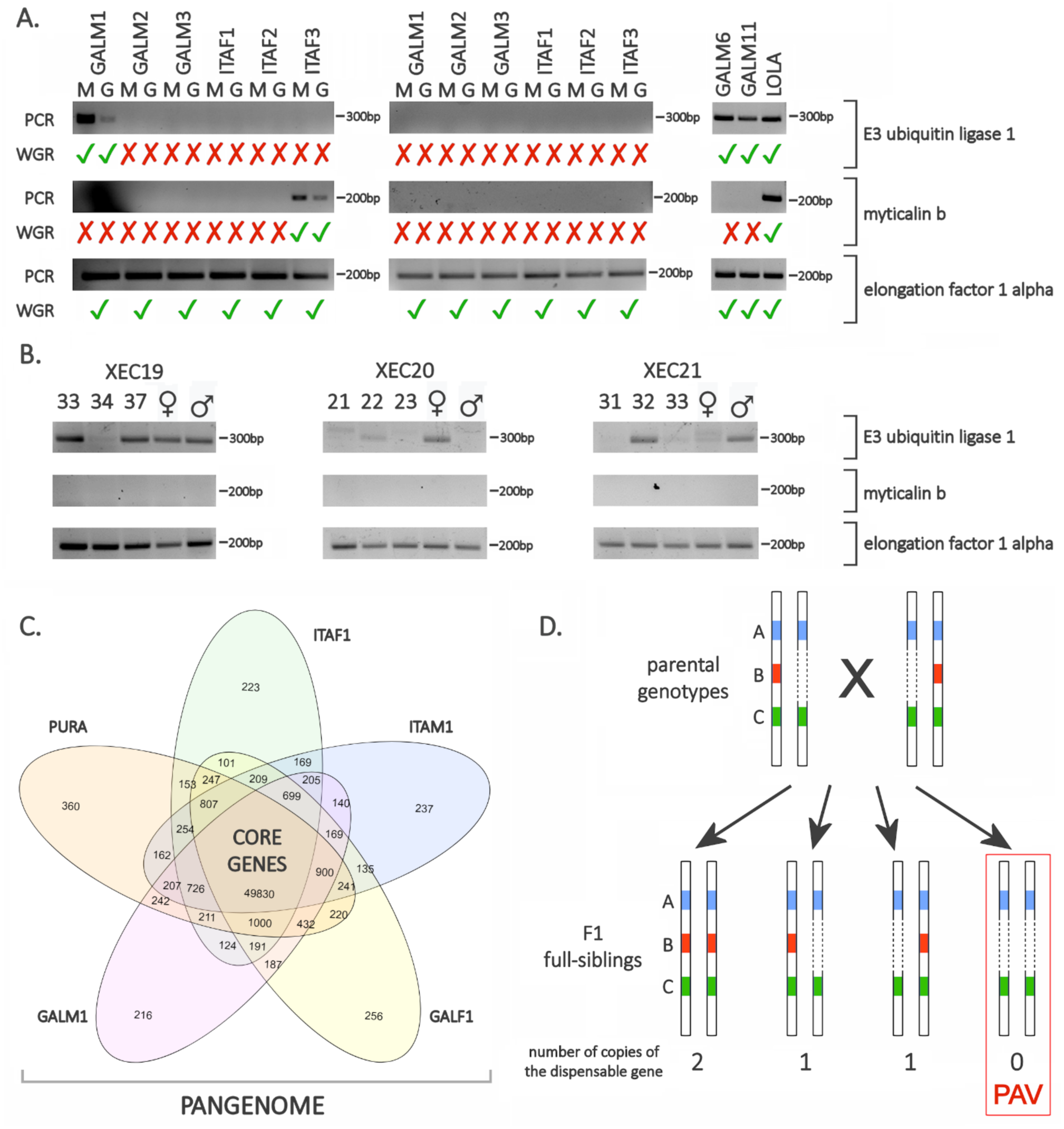
Panel A: validation of the presence/absence variation phenomenon by PCR, carried out on the genomic DNA extracted from the mantle (M) or gills (G) of the 14 mussel individuals subjected to whole genome resequencing. In *Lola*, GALM6 and GALM11, genomic DNA was extracted from the mantle tissue only. One *core* gene (elongation factor 1 alpha) and two *dispensable* genes (E3 ubiquitin ligase 1 and myticalin B1) were tested. Tick and cross symbols indicate expectations based on the *in silico* analysis of whole-genome resequencing (WGR) data. Panel B: observation of the presence/absence variation phenomenon by PCR carried out in 3 full-sib mussels obtained from a controlled cross (parents were also tested, and are indicated by ♂ and ♀, respectively). One *core* gene (elongation factor 1 alpha) and two *dispensable* genes (E3 ubiquitin ligase 1 and myticalin B1) were tested. XEC19, XEC20 and XEC21 indicate the three mussel families subjected to this investigation. The original photographs of the agarose gels used for the preparation of this figure and technical details about these experiments are available in **Supplementary Data Note 11**. Panel C: structure of the mussel pangenome, as exemplified by a Venn diagram representing the overlap between the gene sets found in *Pura* (a female individual previously sequenced by Murgarella and colleagues(8)) and in four resequenced genomes, i.e. ITAF1, ITAM1, GALF1 and GALM1 (note that all these genes are preset in *Lola*). Genes shared by all individuals define the *core* genome, whereas genes shared by some, but not all, individuals are *dispensable*. The entire complement of *core* and *dispensable* genes defines the mussel pangenome. Panel D: schematic overview of the possible origin of a gene presence/absence variation phenomenon. A cross between two parents carrying two *core* genes (A and C) and one *dispensable* gene (B) is depicted. In this case, both parents carry a single copy of the *dispensable* gene (i.e. the *dispensable* gene is present in a hemizygous genomic region). Based on Mendelian inheritance, PAV should be observed in 25% of the offspring produced by this cross.

### The mussel pan-genome

A pan-genome contains a set of *core* genes present in all individuals and fundamental for survival, and *dispensable* genes, which are only found in a subset of individuals of the same species and usually cover accessory functions (36). The widespread occurrence of PAV in mussels is most certainly consistent with this definition, and it reveals an “open” pan-genomic architecture, with a high rate of *dispensable* to core *genes*, i.e. ∼1:3 (**Figure 3C**). Pan-genomes are frequently found in viruses, where they are thought to provide an evolutionary advantage by enabling a quick response to selective pressure (20), and in bacteria, where *dispensable genes* cover accessory functions linked with an enhanced ability of particular strains to colonize new ecological niches (18). Although the small size, simple organization and fast gene gain, loss and horizontal transfer rates of bacterial genomes (19, 37) can explain the presence of a large number of *dispensable* genes in these organisms, pan-genomes have been also occasionally reported in plants, fungi and microalgae. In some cultivated crops, *dispensable* genes contribute significantly to intraspecific phenotypic diversity and to the development of agronomic traits (15–17). The pan-genome architecture of the coccolithophore *Emiliania huxley* is thought to have an adaptive value, explaining its cosmopolitan oceanic distribution and ability to thrive in different habitats, due to acquisition of accessory metabolic activities provided by *dispensable* genes (35). Recent studies have also revealed the fundamental role of *dispensable* genes in increasing the pathogenic potential and antimicrobial resistance in fungi (21, 22).

In spite of the growing number of reports of pan-genomes in eukaryotes and the recent extension of such investigations to human populations (32), so far the impact of PAV on the intraspecific diversity of animals has been presumed to be minimal, and sometimes linked to the development of pathologies (38). To the best of our knowledge, PAV has been only marginally explored in bivalves as a phenomenon linked to gene families involved in immune functions, i.e., big defensins in *Crassostrea gigas* (39) and myticalins in *M. galloprovincialis* (40). Therefore, this is the first study to report the widespread occurrence of PAV at a whole genome scale in metazoans, and the very first report of a pan-genome in the animal kingdom. Although 60,338 protein-coding genes are present in *Lola*, the open pan-genome architecture of the mussel genome suggests that its actual size might be much larger, possibly exceeding 70,000 genes (**Supplementary Data Note 19**). This idea is supported by: (i) the very low frequency of some *dispensable* genes in the resequenced individuals; (ii) the large number of expressed protein-coding mRNAs encoded by *dispensable* genes that are absent in *Lola* but present in other resequenced individuals (**Supplementary Data Note 12**); and (iii) the weak correlation between PAV patterns and the geographical origin of the resequenced individuals (**Supplementary Data Note 19**).

### PAV genes may be associated with survival

In spite of their impact on phenotypic diversity, *dispensable* genes in plants are generally subject to more relaxed selective constraints compared to *core* genes, and they often display traits of reduced essentiality, such as recent acquisition, high tissue specificity, poor expression and increased rate of loss-of-function mutations (22, 41). Mussel genes subject to PAV share similar features, i.e., they display lower levels of expression, shorter ORF length and minor gene architecture complexity compared to *core* genes. They are also on average younger, subject to an increased lineage-specific duplication rate and four times more likely to be taxonomically-restricted compared to *core* genes (**Supplementary data Note 16-17**). However, at the same time, mussel *dispensable* genes retain signatures of functionality, including the presence of conserved regulatory elements and lack of significant GC or codon usage bias (**Supplementary Data Note 13-14**).

We identified several mussel gene families significantly more subject to PAV than expected by chance (**Figure 4A, Supplementary Data Note 15**). The functional annotation of the *dispensable* genes revealed an enrichment in functions related to survival, which may be provided by proteins with marked protein- or carbohydrate-binding properties (e.g., pattern recognition receptors like C1qDC proteins, FReDs and Ig domain-containing proteins), involved in apoptosis pathways (e.g. DEATH and BIR) or playing a role in immune signaling (e.g., Interferon-inducible and IMAP GTPases). Mussel antimicrobial peptides (AMPs) are also subject to massive gene PAV. Although several dozen different sequence variants were identified for each AMP family in the resequenced genomes, each individual mussel possesses a unique combination of a distinct, but small, number of variants, with very little overlap with other mussels from the same population. For example, each individual only possesses, on average, nine out of the 63 myticin variants identified (**Figure 4B, Supplementary Data Note 18**). In light of this observation, PAV emerges as the main factor in determining the high intraspecific sequence diversity observed both at the mRNA and protein level in mussel AMPs (39, 40). On the other hand, other gene families were significantly under-represented among *dispensable* genes. Notably, these included genes encoding transposable elements (and therefore found in multiple copies in the genome) or with housekeeping functions (e.g. protein kinases and G protein-coupled receptors).

**Figure 4:**
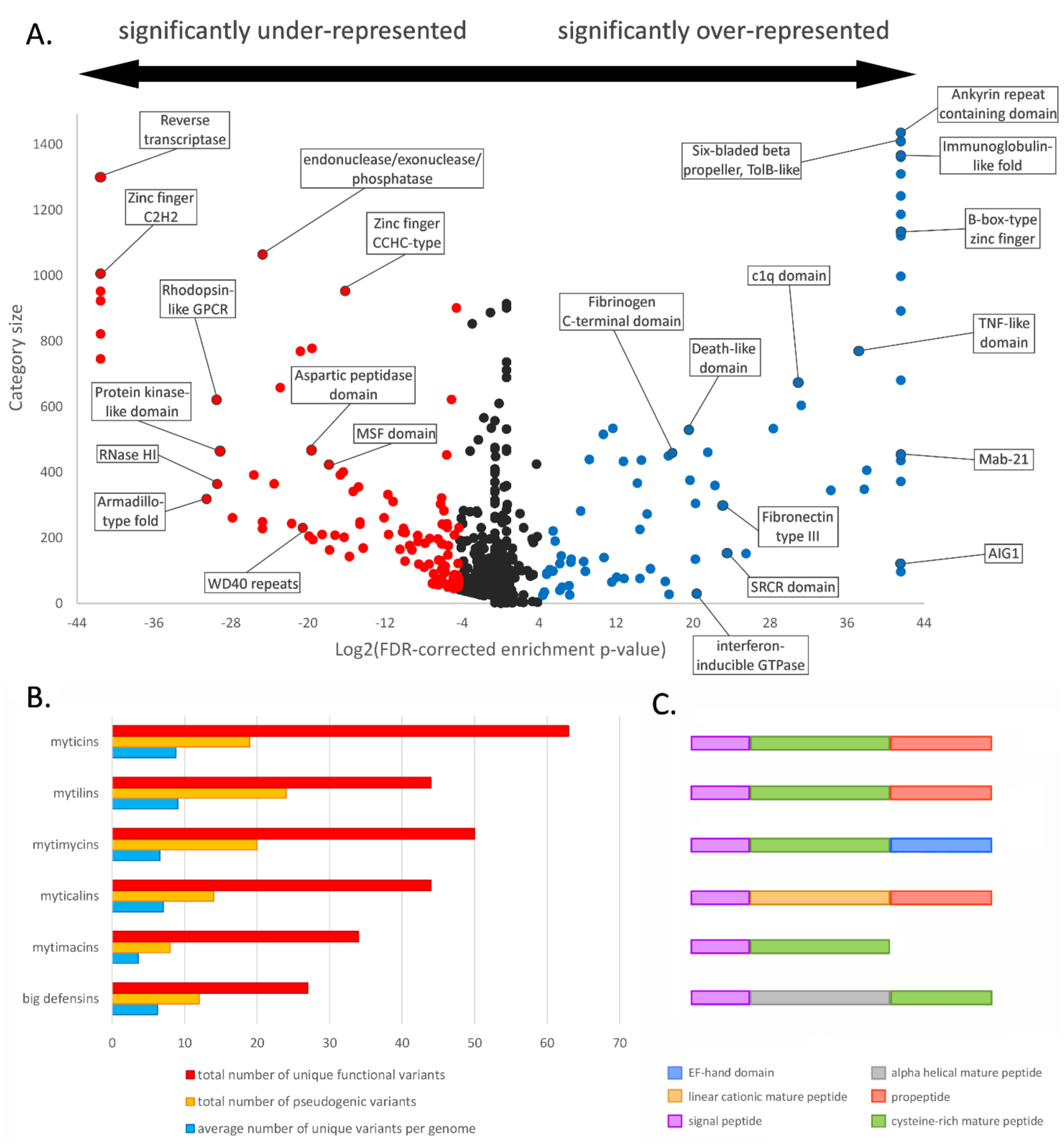
Panel A: correlation between abundance of conserved protein domains (Y axis) and their level of enrichment in the *dispensable* mussel genes (X axis). Each dot represents a conserved protein domain, with specific examples highlighted within boxes. Domains shown in red and blue are under-and over-represented, respectively, in the *dispensable* gene set compared to the *core* genome. Panel B: total number of unique potentially functional sequence variants and pseudogenes, and average number of unique functional variants per genome for the six families of antimicrobial peptides analyzed (myticins, mytilins, mytimycins, myticalins, mytimacins and big defensins). Panel C: schematic organization of the precursor proteins of the six AMP families displayed in panel B.

### Origins and maintenance of PAV in mussels

The *Mytilus* genus has a complex evolutionary history, characterized by extensive gene flow among congeneric species, a process which is still ongoing in mosaic hybrid zones (42–44). However, the analysis of nuclear and mitochondrial genetic markers ruled out the possibility that our resequenced individuals were hybrids between *M. galloprovincialis* and other *Mytilus* species (**Supplementary Data Note 7**), which suggest that *dispensable* genes are unlikely to be recently introgressed allelic variants that cannot be mapped to the reference genome due to their sequence divergence (**Supplementary Data Note 9**). While genetic admixture among contemporary mussel species cannot explain the mussel pan-genome architecture, the role of ancient hybridization and homologous recombination between ancestral *Mytilus* species remains to be investigated, as similar processes have been identified as the key drivers of PAV in plants (45).

Regardless of the origins of mussel *dispensable* genes, their presence in zero, one or two copies in the mussel genome suggests that the PAV phenomenon might be strictly dependent on the matching between paternal and maternal chromosomes during the fertilization process, and that *dispensable* genes might have Mendelian inheritance (**Figure 3D**). To test this hypothesis, we carried out a controlled cross with individuals showing PAV at the E3 ubiquitin ligase 1 gene, resulting in full-sibs showing F1 proportions fully compatible with a Mendelian pattern (**Figure 3B**).

Our finding that a large fraction of the mussel genome is in a single-copy state is congruent with the presence of polymorphic chromosomes (46) and with the significant intra-individual and inter-population variation in nuclear DNA content reported in previous cytogenetic studies (47). The very high amount of intraspecific genomic diversity revealed in our study may come at the cost of interfering with conventional homologous chromosome pairing, recombination and segregation during meiosis.

We observed highly skewed coverage profiles in the sequencing libraries from the gonadal tissue of some (but not all) male mussels, regardless of the stage of sexual maturation. These results, confirmed by a second independent round of resequencing, were not obtained in non-reproductive tissues (i.e., gills) and in female individuals (**Figure 2A**). We suspect that this observation may be the result of a massive presence of aneuploid gametes, potentially generated by an aberrant meiotic process linked with the high structural divergence between homologous chromosomes (**Supplementary Data Note 20**). Several studies have reported the presence of strong genetic barriers in *Mytilus*, acting both between and within species. Although intrinsic postzygotic selection has been invoked as one of the most likely mechanisms underpinning the preservation of mosaic hybrid zones (44, 48), the nature of this process still remains to be elucidated. Here we postulate that the reduced fertility of the offspring produced by individuals carrying “structurally incompatible” chromosomes may be key for explaining post-zygotic selection and the maintenance of the pan-genome architecture in mussels.

Whether the pan-genomic architecture of the mussel genome provides a selective advantage at the population level is a fundamental question. In other organisms, *dispensable* genes can be beneficial, for example enhancing the ability to acquire novel metabolic pathways (e.g., bacteria) (18), to migrate to new ecological niches (e.g., bacteria and unicellular algae) (35) or to enhance the ability to interact with a host (e.g., viruses and fungal pathogens) (20–22). In our opinion, the large over-representation of genes involved in the response to stress and survival in the variable fraction of the mussel pan-genome (**Supplementary Data Note 15**) and the impact of PAV on the molecular diversification of AMPs (**Figure 4B, Supplementary Data Note 18**) strongly suggest an adaptive role for the pan-genomic architecture. This would be consistent with the benefits provided by the development of a complex arsenal of immune molecules in sessile species characterized by high population densities such as mussels, where the spread of pathogens can be very efficient (49, 50). We can speculate that the accessory functions provided by the 15,000 *dispensable* mussel genes might underpin an improved ability to adapt to challenging and varying environmental conditions, resulting in the cosmopolitan distribution and high invasiveness potential of this species (2, 4). Nevertheless, a large number of genes subject to PAV pertain to expanded, taxonomically-restricted gene families with unknown function (**Supplementary Data Note 16-17**). Therefore, the adaptive benefits of PAV might extend well beyond immunity and survival, with a potential impact on multiple aspects of mussel biology. Indeed, specific functional studies would be necessary to confirm these hypotheses.

## Conclusions

We provide, for the first time, significant evidence in support of a pan-genome in the animal kingdom. The unusual structure of the mussel genome is the result of massive levels of gene PAV along single-copy regions, which contain several thousand protein-coding genes. The enrichment of *dispensable* genes in functions linked to resilience to stress and immune response warrant further investigation on the possible links between massive PAV and the evolutionary success of mussels, exemplified by the cosmopolitan distribution of this species in temperate marine coastal waters. Most likely, extensive PAV might be found in other cosmopolitan marine invertebrates characterized by broadcast spawning, very large effective population size and subject to similar environmental pressures.

## Authors contribution

BN, CC, DP, and AF planned and granted the funding to start the project. CC, DP, AF, TA and MGu designed the sequencing strategy. RM extracted the genomic DNA from Lola and all resequenced individuals. MGu performed the Illumina sequencing of Lola. FC, LF, AC and TA performed the genome assembly. JGG, AV and TA performed the genome annotation. MM and CC carried out preliminary analyses of the genome of Lola and provided their inputs for the integration of genomic data from Pura in the present manuscript. MGe, AP and AF planned the whole genome resequencing approach from additional individuals. PV and UR provided mussels from the Adriatic Sea. AF, BN, AP and MGe managed and analyzed sequencing data. FC and TA performed the single-nucleotide variation analysis. PB generated the full-sib mussel families. RM validated PAV from the sequenced individuals and full-sib mussel families. TG, MNO and DP carried out the phylogenetic analyses and studied the evolutionary origin of dispensable genes. MGe and UR analyzed RNA-sequencing data. MGe, AP, PV, UR, RM, AF and BN performed the analysis of genes encoding antimicrobial peptides. MGe, AF, and DP wrote the manuscript with inputs from the other authors. All authors contributed to the writing of the supplementary data notes and to the preparation of supplementary tables and figures. All the authors read and approved the manuscript. DP and AF supervised the whole study.

## Materials & correspondence

All the sequencing data obtained in this work, as well as the genome assembly and the annotation are available in the European Nucleotide Archive (ENA) under the project ID PRJEB24883.

A genome browser and a blast server for this genome can be accessed in our local server (http://denovo.cnag.cat/genomes/mussel/).

The mussel phylome is available for download or browsing at PhylomeDB (http://www.phylomedb.org).

## Acknowledgements

This work was conducted with the support of the projects AGL2011-14507-E, AGL2015-65705-R, RTI2018-095997-B-I00 (Ministerio de Ciencia, Innovación y Universidades, Spain) and INCITE 10PXIB402096PR, IN607B 2016/12 (Consellería de Economía, Emprego e Industria - GAIN, Xunta de Galicia). Antonio Figueras, Beatriz Novoa, Rebeca Moreira, Alberto Pallavicini, Marco Gerdol, Paola Venier and Umberto Rosani are supported by the European Union’s Horizon 2020 research and innovation programme under grant agreement No. 678589. David Posada is supported by the European Research Council, the Spanish Ministry of Economy and Competitiveness and Xunta de Galicia. We would like to thank the following members of the CNAG Data Analysis Team for their help evaluating intermediate assemblies and their useful comments on Variant Calling: Sophia Derdak, Marcos Fernández, Steve Laurie, Jordi Morata and Raúl Tonda. We would like to thank Samuele Greco (Department of Life Sciences, university of Trieste) for his technical support in the bioinformatics analysis of whole-genome resequencing data. We are grateful to Edoardo Turolla and the staff from Istituto Delta (Goro, Italy) for the support provided in mussel sampling. We would like to thank María Gasset for critically reading the manuscript and providing suggestions for its improvement.

